# Brown adipose tissue thermogenesis rhythms are driven by the SCN independent of adipocyte clocks

**DOI:** 10.1101/2024.10.28.620609

**Authors:** Georgios K. Paschos, Ronan Lordan, Taylor Hollingsworth, Damien Lekkas, Sean Kelch, Emanuele Loro, Ioannis Verginadis, Tejvir Khurana, Arjun Sengupta, Aalim Weljie, Garret A. FitzGerald

## Abstract

Circadian misalignment has been associated with obesity both in rodents and humans. Brown adipose tissue (BAT) thermogenesis contributes to energy expenditure and can be activated in adults to reduce body weight. Although previous studies suggest control of BAT thermogenesis by the circadian clock, the site and mechanisms of regulation remain unclear. We used mice with genetic disruption of the circadian clock in the suprachiasmatic nucleus (SCN) and peripheral tissues to delineate their role in BAT thermogenesis. Global post-natal deletion of *Bmal1* in adult mice (*Bmal1^-/-^*) abolishes the rhythms of interscapular BAT temperature, a measure of thermogenesis, while normal locomotor activity rhythms are maintained under a regular 12h light-12h dark schedule. Activation of thermogenesis either by exposure to cold or adrenergic stimulation of BAT displays a diurnal rhythm with higher activation during the active period. Both the rhythm and the magnitude of the thermogenic response is preserved in *Bmal1^-/-^* mice. In contrast to mice with global deletion of *Bmal1*, mice with brown adipocyte (Ucp1-*Bmal1*^-/-^) or brown and white adipocyte (Ad-*Bmal1*^-/-^) deletion of *Bmal1* show intact rhythms of BAT thermogenic activity. The capacity of Ucp1-*Bmal1*^-/-^ mice to activate thermogenesis in response to exposure to cold is identical to WT mice, independent of time of stimulation. Circadian rhythmicity of interscapular BAT temperature is lost in mice with SCN deletion of Bmal1 (SCN-*Bmal1*^-/-^), indicating control of BAT thermogenesis rhythms by the SCN. Control mice exhibit rhythmic BAT glucose and fatty acid uptake - a rhythm that is not recapitulated in *Bmal1*^-/-^ and SCN-*Bmal1*^-/-^ mice but is present in Ucp1-*Bmal1*^-/-^ and Ad-*Bmal1*^-/-^ mice. BAT cAMP and phosphorylated hormone-sensitive lipase (pHSL) is reduced during the active period in *Bmal1*^-/-^ and SCN-*Bmal1*^-/-^ mice consistent with reduced sympathetic tone. Furthermore, sympathetic denervation of BAT ablates BAT temperature rhythms in WT mice. Taken together, our findings suggest that the SCN drives rhythms of BAT thermogenesis through adipocyte clock-independent, sympathetic signaling to the BAT.

## Introduction

Many aspects of modern lifestyle, such as aberrant daily schedules, shift work and sleep disorders create mistiming between the rhythmic environmental daily changes in light availability and temperature, and the endogenous circadian rhythms that are meant to anticipate and prepare us for environmental changes (Boivin et al., 2022). Desynchrony between environmental and physiological rhythms increases the risk for obesity and diabetes (Jaiswal et al., 2020; Liu et al., 2018; Pan et al., 2011; Roenneberg et al., 2012; Sun et al., 2018). The number of years in rotating shift work and the number of night shifts per week are associated with an increased odds ratio for being obese in both men and women (Liu et al., 2018; Pan et al., 2011; Sun et al., 2018). Social jetlag, the difference in sleep midpoint between weekdays and the weekend, a measure of misalignment between the environment and the endogenous clock, has been associated positively with increased body mass index (Roenneberg et al., 2012). Short sleep duration and sleep variability, an index of poor sleep quality, are associated with increased body weight(Jaiswal et al., 2020).

Endotherms generate heat to maintain core body temperature when challenged by low environmental temperature. Brown adipose tissue (BAT) is a major site of non-shivering thermogenesis characterized by high glucose and fatty acid uptake and increased capacity for oxidative phosphorylation in mitochondria (Chouchani et al., 2019). Maximal activation of BAT thermogenesis can increase whole-body energy expenditure by 40%-80% in humans (Ouellet et al., 2012), rendering it a target for strategies aimed at augmenting metabolic rate to counteract obesity and related metabolic disorders. Thermogenesis is subject to strict regulation as it is crucial for the survival of endothermic mammals during cold exposure (Cannon and Nedergaard, 2004; Chouchani et al., 2019). Although the circadian clock has been proposed as one of the regulators of thermogenesis (Chappuis et al., 2013; Gerhart-Hines et al., 2013) the precise mechanisms by which this occurs and the tissues from which this regulation originates are unknown. In this study, we identify the suprachiasmatic nucleus (SCN) as a regulator of daily rhythms in BAT thermogenesis by imposition of rhythmic sympathetic signaling.

## Results

### Disruption of the circadian clock attenuates rhythms of BAT thermogenesis

To assess diurnal rhythmicity in BAT thermogenesis, we applied telemetry to monitor continuously BAT temperature. A telemetry probe was implanted in the interscapular area above the BAT and mice were free to move undisturbed in their home cage. C57BL/6 wild-type mice housed at a room temperature of 22°C exhibited diurnal oscillations of BAT temperature with a peak temperature during the early night, the active phase of the daily cycle (Figure 1A). Mice housed at 22°C need to produce heat to maintain body temperature and as a result they spontaneously activate thermogenesis (Cannon and Nedergaard, 2011). We measured BAT temperature in mice housed at room temperature of 29°C (Figure 1B), to avoid the effect of spontaneous activation on rhythms of BAT thermogenesis. This is a thermoneutral environmental temperature at which the maintenance of mouse body temperature does not require production of heat (Cannon and Nedergaard, 2011). As in mice housed at 22°C, mice at thermoneutral environmental conditions show rhythms of BAT temperature with a peak during the early night (Figure 1A, B). The average BAT temperature recorded in mice housed at thermoneutral environmental temperature was approximately 0.5°C higher compared to the BAT temperature recorded at 22°C. This resulted from the effect of the increase in environmental temperature on the temperature recorded at the probe and does not indicate an increase in BAT temperature at 29°C.

To evaluate the effect of the circadian clock on BAT temperature rhythms we used mice with an inducible universal deletion of the core clock gene *Bmal1* (*Bmal1^-/-^* mice). The absence of *Bmal1* results in loss of functional circadian clocks (McDearmon et al., 2006). Using this mouse model, we induced the deletion of *Bmal1* in adult mice to avoid the developmental defects associated with the lack of BMAL1 (Yang et al., 2016). *Bmal1^-/-^* mice showed severely attenuated BAT temperature rhythms under room temperature of both 22°C and 29°C (Figure 1A, B), while the mice maintained rhythms of locomotor activity (Figure 1C, D). *Bmal1^-/-^*mice retain rhythmic behavior under light: dark conditions despite lacking functional circadian clocks due to the light masking phenomenon (Yang et al., 2016). Our findings indicate that attenuation of BAT temperature rhythms in *Bmal1^-/-^* mice results from the loss of circadian clock function rather than the attenuation of behavioral and activity rhythms.

To exclude the possibility that the deletion of *Bmal1* affected BAT temperature through changes in insulation of *Bmal1^-/-^* mice we measured heat emanated from *Bmal1^-/-^*mice using infra-red camera thermography. The heat detected from both the interscapular region and the rest of the body surface of *Bmal1^-/-^*mice was comparable to the heat emanated from control mice (Figure 1E, F and S1A). In agreement with intact skin insulation, skin morphology was undisturbed in *Bmal1^-/-^* mice. Dermal thickness, thickness of subcutaneous adipose tissue and the number of hair follicles were comparable to control mice (Figure 1G, H and S1B, S1C).

### Maximal activation of BAT thermogenesis does not depend on functional circadian clocks

After excluding heat loss through the skin as the cause of the attenuated BAT temperature rhythms observed in *Bmal1^-/-^* mice, we then investigated the capacity of BAT from *Bmal1^-/-^* mice to produce heat. To stimulate BAT thermogenesis, we exposed 12-14 week-old male mice to a cold environment. Exposure to cold is the physiological stimulus for BAT thermogenesis activation (Cannon and Nedergaard, 2011). Mice housed at 22°C were exposed to 4°C at ZT12 for 3 hours to get acclimatized to cold exposure. Following that, mice were housed at 29°C for 72 hours before they were exposed to 4°C again at either ZT4 or ZT16. Energy expenditure increases rapidly in response to exposure to cold and the increase is mainly due to BAT thermogenesis and muscle shivering in animals adapted to activate BAT thermogenesis (Cannon and Nedergaard, 2011). This was the case for the animals in our experiment. We measured whole body oxygen consumption as a measure of energy expenditure and found that wild-type mice exposed to cold increased oxygen consumption by approximately 3-fold (Figure 2A, B). The increase was higher when cold exposure took place during the night (ZT16), consistent with increased BAT thermogenic activity during the night compared to daytime. *Bmal1^-/-^*mice showed an increase in oxygen consumption in response to cold exposure that was equivalent to wild-type mice (Figure 2A, B). BAT temperature of *Bmal1^-/-^*mice in response to cold exposure was not different from control animals, irrespective of time of day (Figure 2C, D). Mice exhibited only a small increase in locomotor activity and food intake in response to cold exposure because of habituation by virtue of their previous exposure to cold (Figure 2E-H). Importantly, locomotor activity and food intake of *Bmal1^-/-^* mice in response to cold exposure didn’t differ in comparison to control mice (Figure 2E-H).

Mice activate shivering thermogenesis in response to exposure to cold which contributes to the increase in energy expenditure observed under cold exposure. To exclude the possibility that the contribution of shivering in the increase of energy expenditure in response to cold exposure was different in *Bmal1^-/-^* mice, we compared shivering in response to cold between *Bmal1^-/-^* and control mice. Shivering increased in response to cold exposure to the same extend in *Bmal1^-/-^* and control mice (Figure 2I and S2A-C).

Notably, UCP1 protein abundance in the BAT of *Bmal1^-/-^*mice exposed to cold for 4 hours was higher compared to control mice (Figure 2J). The reduction in the size of BAT lipid droplets after cold exposure was not different between *Bmal1^-/-^* and control mice (Figure 2K). BAT mass and the size of lipid droplets was comparable between the two groups (Figure 2L, S2D).

Further to evaluate the thermogenic capacity of *Bmal1^-/-^*mice, we measured oxygen consumption in response to norepinephrine mimicking sympathetic activation of BAT thermogenesis. To avoid the confounding contributions of thermoregulatory thermogenesis, locomotor activity and food intake to energy expenditure, we measured oxygen consumption in anesthetized animals maintained at thermoneutrality. The increase in oxygen consumption observed in *Bmal1^-/-^* mice was comparable to control mice irrespective of time of norepinephrine administration (Figure 2M, N). More specifically, we repeated the experiment with administration of the fat-specific β3 adrenergic receptor agonist CL316,243. The increase in oxygen consumption it evoked was not different between *Bmal1^-/-^*and control mice irrespective of time of administration (Figure 2O, P). Taken together, our findings strongly support intact BAT thermogenic capacity in *Bmal1^-/-^* mice. This suggests that the attenuated BAT temperature rhythms observed in *Bmal1^-/-^*mice are not the result of a defect in BAT thermogenic capacity.

Next, we tested whether *Bmal1^-/-^* mice will maintain BAT thermogenic activity after prolonged exposure to cold. *Bmal1^-/-^*mice previously housed at 22°C were acclimatized to 29°C for 2 weeks and then exposed to 4°C for 1 week. Gross morphology of BAT from *Bmal1^-/-^* mice appeared normal (Figure S2E) and BAT weight was not different from control mice at thermoneutrality or after 1 week of exposure to cold (Figure S2F). Histological analysis of BAT from *Bmal1^-/-^*mice showed no difference in the size or number of lipid droplets compared to control mice (Figure S2J). The morphology of BAT mitochondria from mice exposed to cold were indistinguishable between *Bmal1^-/-^* and control mice (Figure S2K).

### BAT thermogenesis rhythms are independent of the circadian clock in brown adipocytes

The attenuated rhythms of BAT temperature in *Bmal1^-/-^*mice support the control of BAT thermogenic activity rhythms by the circadian clock. Since circadian clocks in peripheral tissues often drive the rhythmicity in physiological processes integral to the physiology of the tissue, we tested whether the circadian clock in brown adipocytes drives BAT thermogenesis rhythms. To disrupt the circadian clock, we deleted *Bmal1* specifically in brown adipocytes (Ucp1-*Bmal1*^-/-^) (Figure 3A) and measured BAT temperature. Telemetry monitoring showed intact BAT temperature rhythms under room temperature and thermoneutrality (Figure 3B, C) while as expected, the locomotor activity rhythms of the Ucp1-*Bmal1*^-/-^ mice was unaffected (Figure 3D, E). Relative to littermate controls, Ucp1-*Bmal1*^-/-^ mice exhibited similar body weights, basal energy expenditure, and food intake (Figure S3A-F). Consistent with intact BAT thermogenesis capacity, the Ucp1-*Bmal1*^-/-^ mice exhibited an increase in energy expenditure in response to cold exposure comparable to control mice independent of time of cold exposure (Figure 3F, G). BAT temperature in response to cold was not different between Ucp1-*Bmal1*^-/-^ and control mice (Figure 3H, I). Mice exhibited a minimal increase in locomotor activity and food intake in response to cold exposure and no difference was observed between Ucp1-*Bmal1*^-/-^ and control mice (Figure 3J-M). Gross morphology of BAT from Ucp1-*Bmal1*^-/-^ mice appeared normal (Figure 3N) and BAT weight was not different to control mice at 22°C or after 4 hours of exposure to cold (Figure 2O). UCP1 was higher in BAT from Ucp1-*Bmal1*^-/-^ mice housed at 22°C but not after 4 hours of exposure to cold (Figure 3A). Histological analysis of BAT from Ucp1-*Bmal1*^-/-^ mice showed no difference in the size or number of lipid droplets compared to control mice (Figure 3P and S3G).

### BAT thermogenesis rhythms are independent of peripheral circadian clocks

Activation of BAT thermogenesis increases triglyceride lipolysis both in brown and white adipose tissue (WAT) (Shin et al., 2017). Non-esterified fatty acids from WAT fuel uncoupled respiration in BAT for thermogenesis (Schreiber et al., 2017). To test whether disruption of the white adipocyte circadian clock disrupts BAT thermogenesis rhythms, we generated mice with a deletion of *Bmal1* in both white and brown adipocytes (Ad-*Bmal1*^-/-^). We observed no difference in BAT temperature rhythms of Ad-*Bmal1*^-/-^ compared to control mice under room temperature and thermoneutrality (Figure S4A, B). BAT resident macrophages remove oxidatively damaged mitochondrial fragments released from brown adipocytes to support the BAT thermogenic response to cold (Rosina et al., 2022). To test the role of the circadian clock in adipose-resident macrophages in BAT thermogenesis rhythms, we deleted *Bmal1* in myeloid cells that include BAT and WAT resident macrophages and measured BAT temperature. Mice without functional adipose macrophage circadian clocks showed normal rhythms of BAT thermogenesis (Figure S4C, D). Finally, rhythms in BAT temperature were preserved in mice with a deletion of *Bmal1* in sympathetic nerves (Figure S4E, F). Collectively, our findings do not support a role of peripheral clocks in cells involved in BAT thermogenesis in the control of BAT thermogenesis rhythms.

### BAT thermogenesis rhythms are driven by the SCN

With no evidence of control by peripheral circadian clocks, we hypothesized that BAT thermogenic activity rhythms are controlled by the SCN. To test our hypothesis, we generated tissue specific *Bmal1* knockout mice to ablate the circadian clock in the SCN, by crossing *Bmal1*^fx/fx^ mice to Six3-Cre mice (Six3-*Bmal1*^-/-^). As previously reported, Six3-Cre is expressed in ventral anterior hypothalamus including the SCN, the subparaventricular zone but not in the dorsomedial hypothalamus (Bedont et al., 2017). To validate deletion of *Bmal1* in the SCN we measured *Bmal1* expression at ZT22 in punch biopsies of the SCN, arcuate nucleus (ARC), the ventromedial hypothalamus (VMH) and the dorsomedial hypothalamic nucleus (DMH). The dissection method was validated by the expression of markers specific for each hypothalamic area (Figure S5A). Expression of *Bmal1* was markedly reduced in the SCN of Six3-*Bmal1*^-/-^ mice but unaffected in nearby hypothalamic nuclei (Figure 4A). Importantly, no BMAL1 protein was detected in SCN from Six3-*Bmal1*^-/-^ mice (Figure 4B). Telemetry monitoring revealed loss of BAT temperature rhythms in Six3-*Bmal1*^-/-^ mice (Figure 4C, D). The lack of BAT temperature rhythms was independent of environmental temperature or changes in locomotor activity (Figure 4E, F). Further, to support control of BAT thermogenesis rhythms by the SCN we availed of a second mouse without a functional SCN clock. Here deletion of *Bmal1* in the SCN was achieved by expressing Cre recombinase under the control of the Synaptotagmin10 (Syt10) promoter in *Bmal1*^fx/-^ mice (Syt10-*Bmal1*^fx/-^) (Husse et al., 2011). In agreement with SCN control of BAT thermogenesis rhythms, Syt10-*Bmal1*^fx/-^ mice showed attenuated rhythms of BAT temperature compared to controls under both room temperature and thermoneutrality (Figure 4G, H). Locomotor activity of Syt10-*Bmal1*^fx/-^ mice under light: dark conditions was retained (Figure 4I, J).

We next evaluated activation of BAT thermogenesis by assessing BAT uptake of glucose and oleic acid during daytime (ZT4) and nighttime (ZT16). Glucose and fatty acid uptake by the BAT are a measure of BAT thermogenic activity (McNeill et al., 2020). BAT uptake of both glucose and oleic acid increased at ZT16 in control mice as compared to the uptake at ZT4 in agreement with rhythmic activation of BAT thermogenesis and higher thermogenic activity during the night (Figure 4K-N, S5B). Consistent with attenuated BAT thermogenesis rhythms, BAT uptake of glucose and oleic acid at ZT16 didn’t increase in *Bmal1*^-/-^, Syt10-*Bmal1*^fx/-^ and Six3-*Bmal1*^-/-^ mice when compared to the increase in control mice (Figure 4K-N, S5B). In contrast, Ucp1-*Bmal1*^-/-^ and Ad-*Bmal1*^-/-^ mice showed an increase of glucose and oleic acid BAT uptake at ZT16 that was like the increase in control mice (Figure 4K-N, S5B). Glucose and oleic uptake by inguinal WAT, visceral WAT, liver and skeletal muscle was indistinguishable between groups at both ZT4 and ZT16 (Figure 4K-N, S5B). Our finding that BAT glucose and oleic acid uptake increased during the night only in mice with a functional SCN clock is consistent with SCN control of BAT thermogenesis rhythms.

### The SCN drives rhythms in BAT thermogenesis by rhythmic sympathetic signaling to BAT

Sympathetic signaling to BAT is the major physiologic signal controlling activation of BAT thermogenesis. Sympathetic neurons release norepinephrine, which activates the β-adrenergic receptor-cAMP-PKA pathway in adipocytes (Sakers et al., 2022). To evaluate whether the SCN drives rhythms of BAT thermogenesis by imposing rhythmic sympathetic signaling to BAT we measured cAMP in BAT at ZT4 and ZT16. BAT cAMP at ZT 16 was reduced in *Bmal1*^-/-^, Syt10-*Bmal1*^fx/-^ and Six3-*Bmal1*^-/-^ mice compared to controls (Figure 5A, S5C), consistent with an attenuated sympathetic signaling rhythm in mice with no functional SCN clock. BAT cAMP was unaffected in Ucp1-*Bmal1*^-/-^ and Ad-*Bmal1*^-/-^ mice (Figure 5A, S5C). Next, we measured phosphorylated hormone sensitive lipase (pHSL), a member of the signaling cascade of the β3 adrenergic receptor to investigate sympathetic signaling to BAT. BAT pHSL was reduced in *Bmal1*^-/-^ mice at ZT16 (Figure 5B) consistent with loss of diurnal rhythms of sympathetic signaling to BAT when the SCN clock is disrupted. Finally, we evaluated rhythms of BAT temperature in mice with sympathectomized BAT to test further our hypothesis that SCN drives rhythms of BAT thermogenesis via rhythmic sympathetic signaling to BAT. Bilateral surgical sympathetic denervation of BAT in C57BL/6 wild-type mice resulted in loss of BAT temperature rhythms both at room temperature and thermoneutral conditions (Figure 5C). Taken together, our findings support the control of BAT thermogenesis rhythms by the SCN through rhythmic sympathetic signaling to BAT.

## Discussion

The circadian control of BAT thermogenesis has been proposed by studies in animal models with genetic deletion of core clock components (Chappuis et al., 2013; Gerhart-Hines et al., 2013; Li et al., 2013). Global deletion of *Rev-erb*α increased BAT UCP1 and the ability of mice to defend body temperature during cold exposure (Gerhart-Hines et al., 2013). In contrast, *Per2* mutant mice were more susceptible to hypothermia with reduced UCP1 in BAT in response to cold exposure (Chappuis et al., 2013). Furthermore, mice carrying an embryonic global deletion of *Bmal1* were indistinguishable from wild-type mice in maintaining the core body temperature under cold exposure (Li et al., 2013). Mice exposed to constant light have reduced sympathetic input into BAT and decreased glucose and lipid uptake by BAT (Kooijman et al., 2015). Conversely, mice lacking Bmal1 in the ventromedial hypothalamus have enhanced BAT activity through increased β3-adrenergic receptor activation that can be restored by restricted feeding (Orozco-Solis et al., 2016). Similarly, deletion of Bmal1 in astrocytes increased interscapular temperature measured by thermographic imaging in mice (Luengo-Mateos et al., 2023).

In this study, we implemented continuous monitoring of BAT temperature to investigate the effect of circadian clock disruption on BAT thermogenesis. We found that BAT thermogenesis rhythms were attenuated but the capacity of BAT to generate heat after stimulation was preserved in mice without functional circadian clocks. We identify the SCN as the regulator of daily BAT thermogenesis rhythms by driving rhythmic sympathetic stimulation of BAT. The SCN directly innervates BAT among other brain structures (Bamshad et al., 1999). BAT thermogenesis is increased after electrical stimulation of the retino-hypothalamic tract that innervates the SCN, or after glutamate injections directly into the SCN in anesthetized rats (Amir et al., 1989). In support of the role of the sympathetic nervous system in activating BAT thermogenesis, mice lacking β-adrenergic receptors show impaired body temperature regulation during cold exposure (Razzoli et al., 2018; Ueta et al., 2012). BAT sympathectomy abolishes rhythms of BAT temperature in our study highlighting the sympathetic nerves as the route for SCN control of BAT thermogenesis rhythms. A drawback of the BAT sympathectomy model we used is that surgical denervation not only severs the descending SNS innervation to the BAT but also the ascending sensory innervation from BAT to the brain (Vaughan et al., 2014). SCN control of BAT thermogenesis rhythms through rhythmic sympathetic stimulation of BAT appears independent of functional circadian clocks in sympathetic nerves. Mice lacking *Bmal1* in sympathetic nerves maintain rhythms of BAT thermogenesis. Thus, sympathetic nerves without functional circadian clocks can still transmit rhythmic signals originating from the SCN to BAT. Together, our data are consistent with the hypothesis that the SCN dictates thermogenesis rhythms in BAT via sympathetic innervation.

Unexpectedly, the control of BAT thermogenesis is independent of white and brown adipocyte circadian clocks. Our findings are in contrast with the results in the study by Hasan et al. (Hasan et al., 2021) showing a small reduction in interscapular temperature measured by thermal imaging in mice with BAT deletion of Bmal1. Although we cannot exclude the method of measuring BAT temperature or experimental conditions as being responsible for this difference, we fed Ucp1-*Bmal1*^-/-^ and Ad-*Bmal1*^-/-^ mice a HFD for one week to replicate the conditions of this study and did not observe a difference in BAT temperature for either group (Figure S5D, E). However, BAT deletion of Bmal1 did cause a small reduction in daytime core body temperature at 22°C and 4°C but not at thermoneutrality (30°C) (Chang et al., 2018).

Our findings provide one more example of the hierarchical structure of the circadian clock in mammals where the SCN serves as the master clock that synchronizes the rest of body’s physiology to the light cycles of the environment (Mohawk et al., 2012). Peripheral circadian clocks such as in adipocytes (Paschos et al., 2012) and muscle (Ehlen et al., 2017) were found to provide feedback to the SCN to orchestrate physiological rhythms. In the case of BAT thermogenesis rhythms, peripheral adipocyte clocks were dispensable because of the control the SCN exerts on BAT through sympathetic signaling.

Central control of BAT thermogenesis opens the possibility of time dependent approaches to increase BAT thermogenesis and increase whole body energy expenditure. Such approaches to correcting circadian misalignment have the potential to improve glucose and lipid metabolism (Chellappa et al., 2021).

## Experimental Procedures

### Animals

Bmal1^flox/flox^ mice were generated as previously described (Paschos et al., 2012). Bmal1^flox/flox^ mice maintained on a C57BL/6J background were bred to mice with Cre-ER (estrogen receptor) (Hayashi and McMahon, 2002) that render Cre-activity inducible by tamoxifen. 8 weeks old mice were given tamoxifen (i.p. 100 mg/Kg body weight per day) for 5 consecutive days to delete Bmal1 in all cells/tissues (*Bmal1^-/-^*). All experiments were performed one month after the last tamoxifen injection to minimize the possibility of residual drug effects. Bmal1^flox/flox^ mice were bred to Ucp1-Cre mice (Kong et al., 2014) (Jax 024670, provided by E. Rosen), Adipoq-Cre mice (Eguchi et al., 2011) (Jax 028020, provided by E. Rosen), LysM-Cre mice (Clausen et al., 1999) (Jax 004781), Six3-Cre mice (Bedont et al., 2017) (provided by S. Blackshaw) to induce brown adipocyte-specific (Ucp1-*Bmal1*^-/-^), adipocyte-specific (Ad-*Bmal1*^-/-^), myeloid cell-specific (LysM-*Bmal1*^-/-^), SCN-specific (Six-*Bmal1*^-/-^) deletion of Bmal1 respectively. We bred Dbh-3’UTR-IRES-CreERT2 mice (Tsarovina et al., 2010) (Jax 020074) with Bmal1^flox/flox^ mice to generate mice with tamoxifen induced deletion of Bmal1 in sympathetic nerves. 6 weeks old mice were given tamoxifen (i.p. 200 mg/Kg body weight per day) for 5 consecutive days to delete Bmal1 in adrenergic neurons (Dbh-*Bmal1^-/-^*). Mice expressing Cre recombinase under the control of the Synaptotagmin10 (Syt10) promoter in *Bmal1*^fx/-^ mice (Syt10-*Bmal1*^fx/-^) (Husse et al., 2011) (provided by H. Oster) were used as a second mouse carrying a deletion of *Bmal1* in the SCN. Mice were housed at 22°C under a 12:12 hour light: dark cycle with free access to food (LabDiet, 5010) and water unless otherwise stated. All experiments during the dark phase of the daily cycle were performed under dim red light. Mice were exposed to cold at ZT12 for 3 hours prior to exposure to cold environmental temperature to get acclimatized. Mice were returned to 22°C for 3 days and housed at thermoneutrality (29°C) for 72 hours before they were exposed to cold to avoid spontaneous activation of BAT thermogenesis. The mice were then exposed to cold (4°C) for 3 hours at either ZT4 (high BMAL1 transcriptional activity) or ZT16 (low BMAL1 transcriptional activity). Transition from 29°C to 4°C was performed over a period of 1 hour. All experiments used age-matched male littermates. Animals used as controls were Bmal1^flox/flox^ (tamoxifen treated for the *Bmal1^-/-^* and Dbh-*Bmal1^-/-^* mice). Mouse genotypes were assessed by PCR on genomic DNA from tail tips. Studies were performed in accordance with procedures approved by the University of Pennsylvania Institutional Animal Care and Use Committee.

### Measurement of BAT temperature by telemetry

Mice were anesthetized using isoflurane. The dorsal area of mice was shaved to remove hair and a 0.5cm vertical incision was made on the interscapular area in the back of the animal to open the skin and insert a telemetry transmitter (G2 emitter, Comprehensive Lab Animal Monitoring System (Oxymax-CLAMS), Columbus Instruments) above BAT for the measurement of temperature. The transmitter was secured in place by 5-0 suture.

### Indirect Calorimetry

Energy expenditure was determined by indirect calorimetry in open-circuit Oxymax chambers of the Comprehensive Lab Animal Monitoring System (CLAMS; Columbus Instruments). Mice were housed individually and maintained at 22°C under a 12:12 hour light: dark cycle with free access to food (LabDiet, 5010) and water. Locomotor activity, food and water intake was measured in the CLAMS chambers. For the assessment of maximal activation of thermogenesis after adrenergic stimulation mice were housed at 33°C for 6 hours prior. Mice were anesthetized using 50 mg/kg pentobarbital i.p. 20 minutes before the administration of the adrenergic agonist. Pentobarbital was the preferred method of anesthesia as inhalation anesthetics and ketamine/xylazine may inhibit brown adipose tissue thermogenic activity. Mice were injected with 1mg/kg norepinephrine or 1mg/kg CL-316,243 subcutaneously either at ZT4 or at ZT16 and were returned to the cage at 30°C to continue monitoring VO_2_.

### Tissue glucose, oleic acid uptake

Mice were individually housed 29°C for 1 week before the experiment. A mixture of 14C-oleic acid (Oleic Acid, [1-14C]-, NEC317050UC, Perkin Elmer) and fatty acid free and endotoxin free bovine serum albumin (A8806-1G, Sigma) in phosphate buffered saline (33 mg/ml) was prepared and filter-sterilized. A separate mixture of 3H-2-deoxy-D-glucose (1-[1,2-3H(N)]-deoxy-D-glucose; NET549250UC; Perkin Elmer) and sterile phosphate-buffered saline was prepared and the two mixtures were combined for a final concentration of 1 μCi/ml of 14C-oleic acid and 2.5 μCi/ml of 3H-2-deoxyglucose. Mice were administered 200μl of the mixture by tail-vein injection either at ZT4 or ZT16. Mice were anesthetized by 1-4% isoflurane 10 minutes after injection, euthanized by cervical dislocation, and plasma and tissues were collected. Tissues were lysed and homogenized using high-speed shaking with a metallic bead in phosphate buffered saline (TissueLyser II, Qiagen). Scintillation solution (5ml, Ecoscint H, LS-275, National Diagnostics) were added to 300μl of the homogenate and radioactivity was measured as cpm/mg tissue with a multi-purpose scintillation counter (LS 6500, Beckman Coulter). Measurements were corrected to plasma radioactivity levels.

### Infrared thermography

Whole body thermal images of mice were captured using Teledyne (FLIR T-530, FLIR Systems, Inc., USA). The thermal images were captured at a distance of 20cm from the mouse housed at 22°C and images were analyzed using the Quick Report, a thermal software with version 1.2 supported by FLIR T-530 camera (www.flir.com).

### Electromyography (EMG)

The EMG setup consisted of three 29-gage needle electrodes (two recording electrodes held stably 3 mm apart and 3 mm deep, and one reference electrode placed distally), applied transcutaneously and connected to a P55 A.C. differential preamplifier (Grass Instruments). The amplified signal was then acquired and A/D converted with a PowerLab 8/SP (ADInstruments). During the recordings, the anesthetized mouse was positioned on a temperature-controlled metal platform connected to a water bath. Core and BAT temperature were monitored with a RET-3 rectal probe for mice (physitemp) and a PFA-insulated 36-gage T-type thermocouple (Omega) respectively, each connected to a T-Type Thermocouple to Analog Converter (Omega).

Mice were anesthetized with an IP injection of urethane 1.5 - 1.8 g/kg, positioned on the metal platform at 33°C, and the basal EMG, core and BAT temperatures were recorded for 10 minutes. Mice were given topical analgesia and allowed to recover for one day. The following day, mice were singly housed and placed at 4oC for 1 hour, then quickly anesthetized and placed on the platform maintained at 15oC for the recordings. The EMG signal was preamplified 1000X and band-pass filtered (3Hz – 10kHz). Root means square (RMS) of the signal was calculated with the LabChart7 software (ADInstruments). For each condition, 4 minutes traces were analyzed, calculating the average RMS signal over 10 seconds segments.

### Histology

Longitudinal dorsal skin sections and interscapular BAT sections were prepared in 4% paraformaldehyde overnight at 4°C. Tissues were paraffin embedded and 15μm sections were used for H&E staining. Images were acquired with an inverted Nikon Labophot-2 microscope and dermal thickness, subcutaneous adipose tissue thickness, number of hair follicles in skin, the number, size of lipid droplets in BAT were measured using ImageJ software and R coding.

### Electron microscopy

Dissected tissues were immediately fixed in 2% paraformaldehyde, 2.5% glutaraldehyde in 0.1M sodium cacodylate buffer at pH 7.4 for at least 24 h. Samples were washed in wash buffer and post-fixed in 2% osmium tetroxide for 1 h at room temperature. The samples were then rinsed in distilled water before *en bloc* staining with 2% uranyl acetate. Following dehydration through a graded ethanol series, the tissue was infiltrated and embedded in EMbed-812 (Electron Microscopy Sciences, Fort Washington, PA, USA). Thin sections were stained with lead citrate and uranyl acetate. The tissues were imaged using a JEOL 1010 transmission electron microscope fitted with a Hamamatsu digital camera and AMT Advantage NanoSprint500 software.

### Gene expression analysis

Total RNA was isolated from tissues using Qiagen RNeasy Mini Kit. Three to four hypothalamic nuclei punches were pooled together per biological replicate for RNA isolation. Reverse transcription was achieved using a Transcriptor First-Strand cDNA Synthesis Kit (Roche). The sequences of the primer pairs are listed in the Supplementary Methods. Quantitative PCR was done on a ViiA7 Real-Time PCR System (Applied Biosystems) using Power SYBR Green. Levels were normalized against Tbp expression.

### Protein extraction and Western blot analysis

Protein was extracted from tissues using RIPA buffer (Boston BioProducts) containing protease and phosphatase inhibitors (Roche) followed by acetone precipatition to remove lipid contamination (Park et al., 2019). For SCN, three biological replicates were combined. For western blot analyses, 30 mg protein was subjected to SDS-PAGE under reducing conditions, transferred, and blotted with an appropriate antibody. The antibodies used were: BMAL1 (Cell Signaling; 14020s at 1:1000), b-ACTIN (Cell Signaling; 4970 at 1:1000), pHSL (Cell Signaling; 45804S at 1:1000), HSL (Cell Signaling; 4107S at 1:1000), TH (Abcam; ab112 at 1:1000), UCP1 (Abcam; ab10983 at 1:2000), Vinculin-HRP (Cell Signaling; E1E9V XP(R) at 1:1000) and Tubulin (Cell Signaling; 2148S at 1:1000).

### Measurement of cAMP

cAMP was profiled using a method described elsewhere (Malik et al., 2018). Briefly, the data was acquired in a Waters TQ-S micro instrument (Waters Corp. Milford, MA) fitted with Waters Acquity H-class UPLC system. Dried samples were reconstituted in 1:1 water/acetonitrile and 2 ul was injected in a XBridge BEH amide column (2.5 μm × 100 mm × 2.1 mm) in analytical duplicates. Chromatographic condition was as follows – starting gradient with 15% A (95:5 water/acetonitrile, 20 mM ammonium acetate adjusted to pH 9 using ammonium hydroxide) and 85% B (acetonitrile) at 0.15 mL/min, followed by a ramp up to 70%A for 10 minutes of isocratic hold. The column was washed after each injection using 98%A and re-equilibrated to starting conditions for 5 minutes post-injection. Quality control (QC) samples made with pooled analytical samples were injected after every ten injections.

cAMP was detected in positive mode using single reaction monitoring approach (m/z 330.0206 → m/z 135.9886) with 50V cone voltage and 30V collision energy. Acquired raw data was exported and converted into mzmol using Proteowizard msconvert (https://proteowizard.sourceforge.io/) followed by integration in El-Maven (Elucidata, San Francisco, CA). Instrumental drift correction was performed using an R script developed in house, followed by normalization with the tissue weight.

### Statistical analysis

Comparisons between two groups were performed with a two-sample *t* test (parametric when the variables passed normality testing and Mann-Whitney otherwise). Comparisons between more than two groups were performed with one-way analysis of variance (ANOVA) or two-way ANOVA and Holm-Sidak’s post-test. Data are presented as means ± SEM and significance was determined at p< 0.05. Statistical analyses were performed with GraphPad Prism v.10.

## Acknowledgments

We thank the Rodent Metabolic Phenotyping Core (RRID: SCR_022427), supported in part by National Institutes of Health (NIH) grant S10-OD025098, the Cox Institute, and the Institute for Diabetes, Obesity and Metabolism (Perelman School of Medicine, University of Pennsylvania) for performing FLIR thermal imaging. We thank the Electron Microscopy Resource Lab for the electron microscopy. The work was supported by a grant from the NIH (U54TR001878) and during this work GAF held a Merit Award from the American Heart Association. GAF is senior advisor to Calico Laboratories.

---

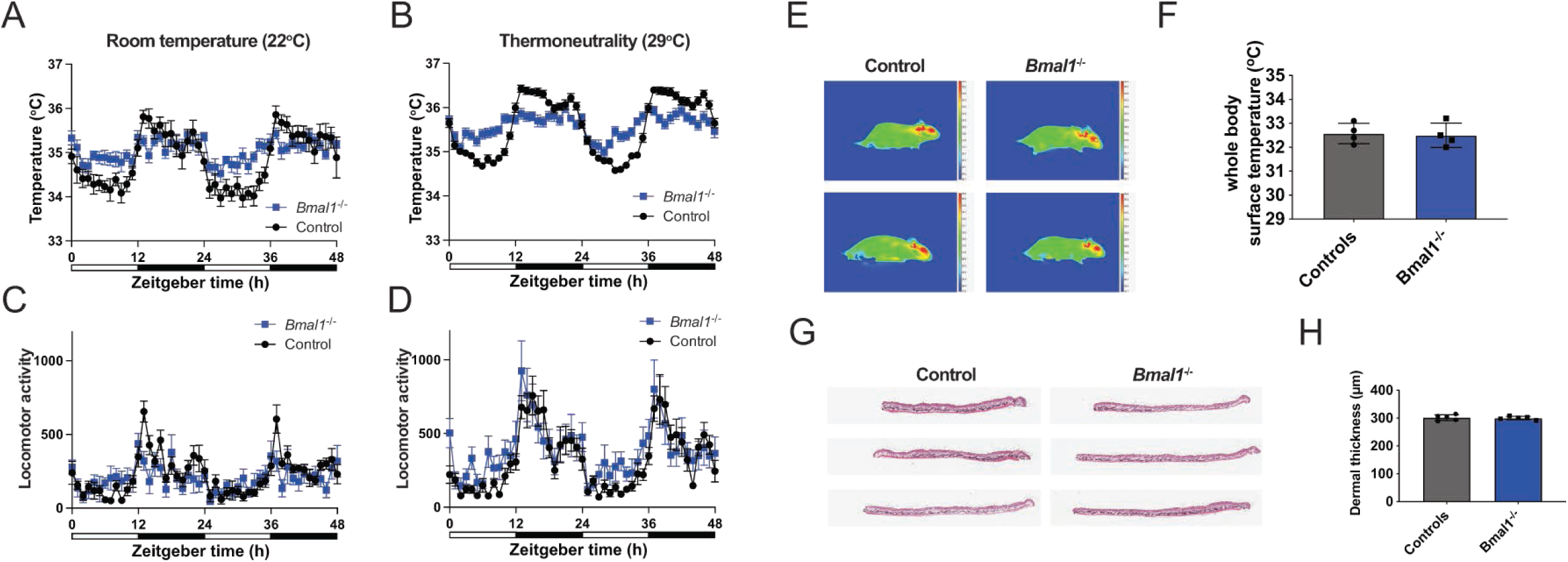

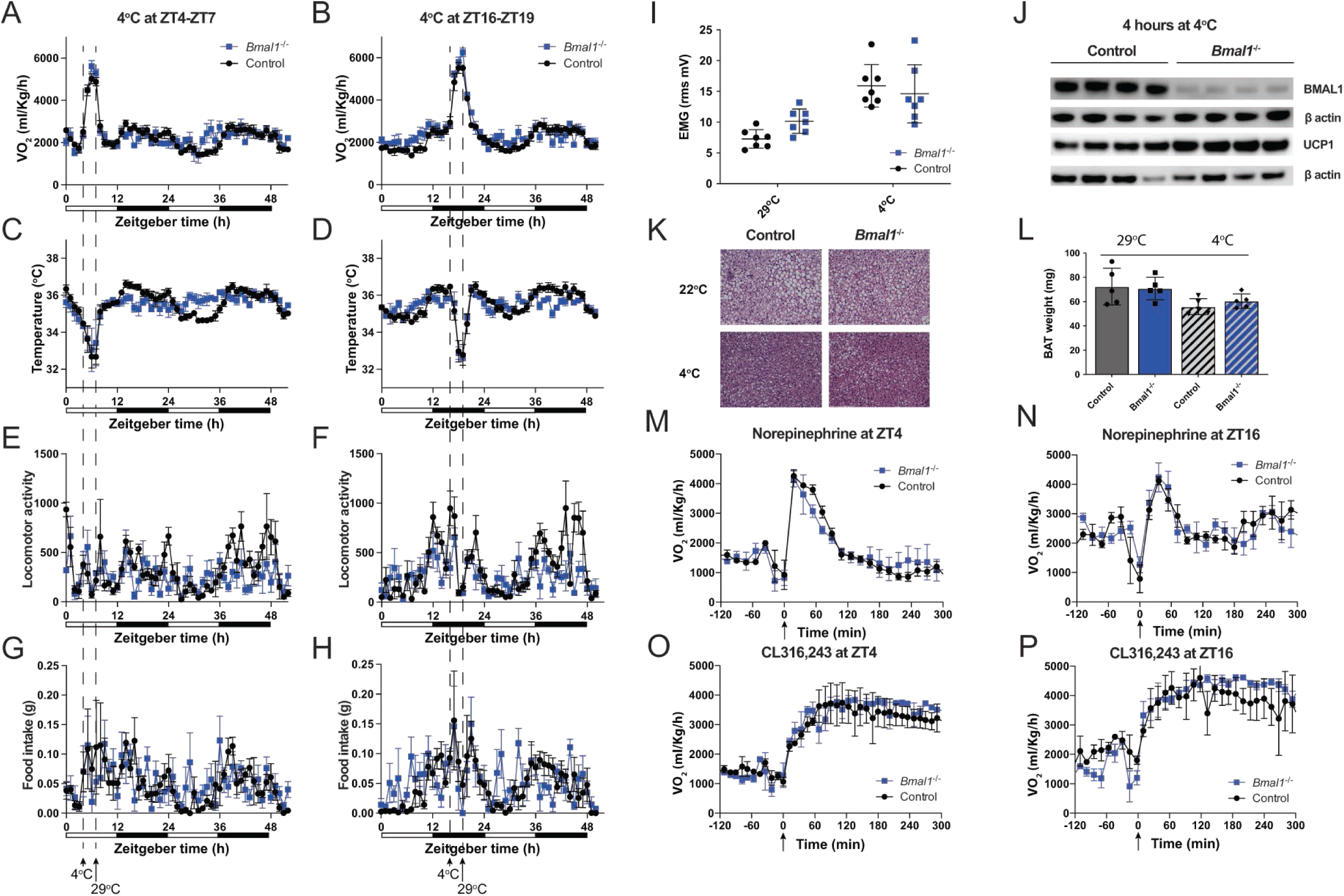

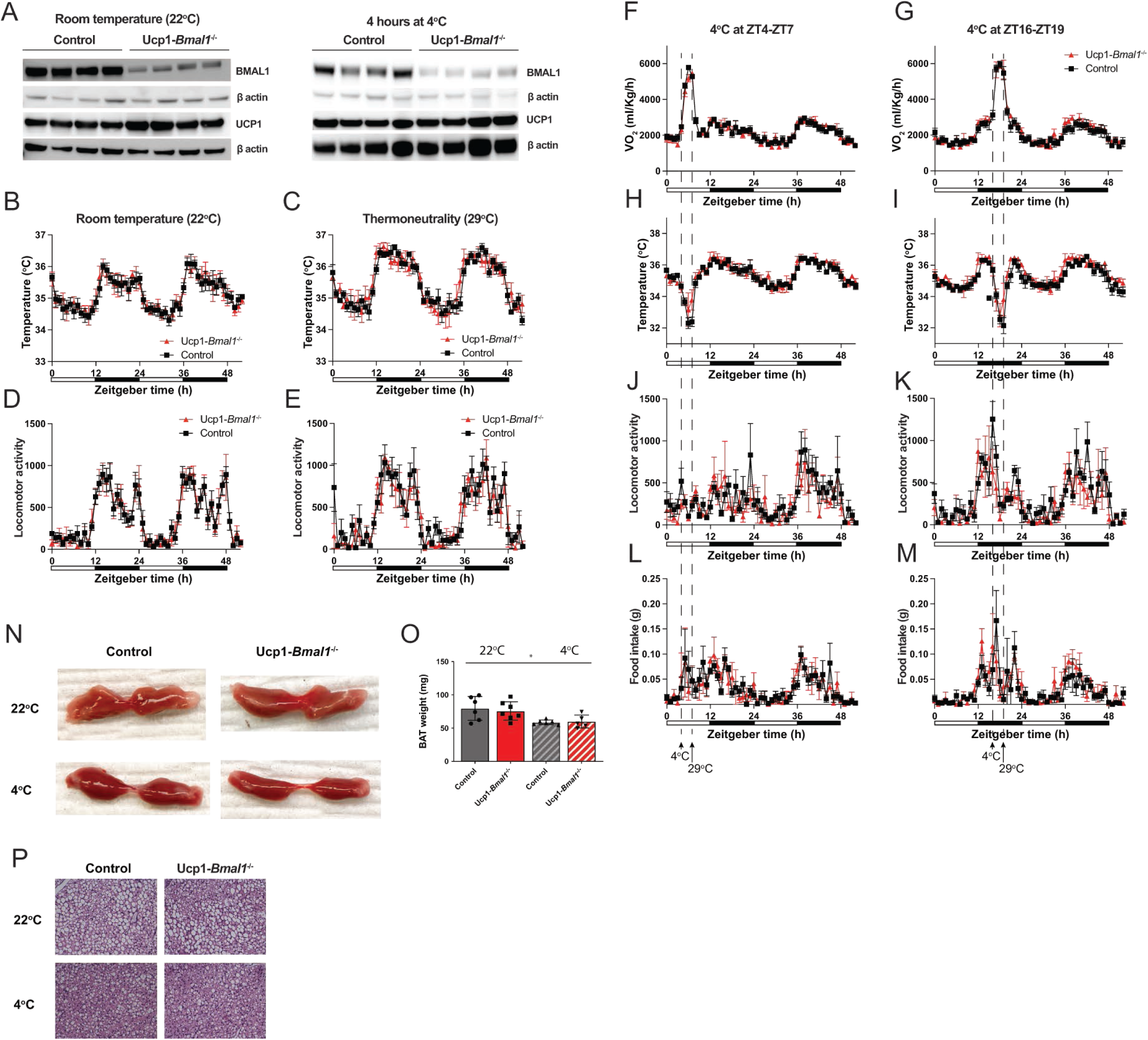

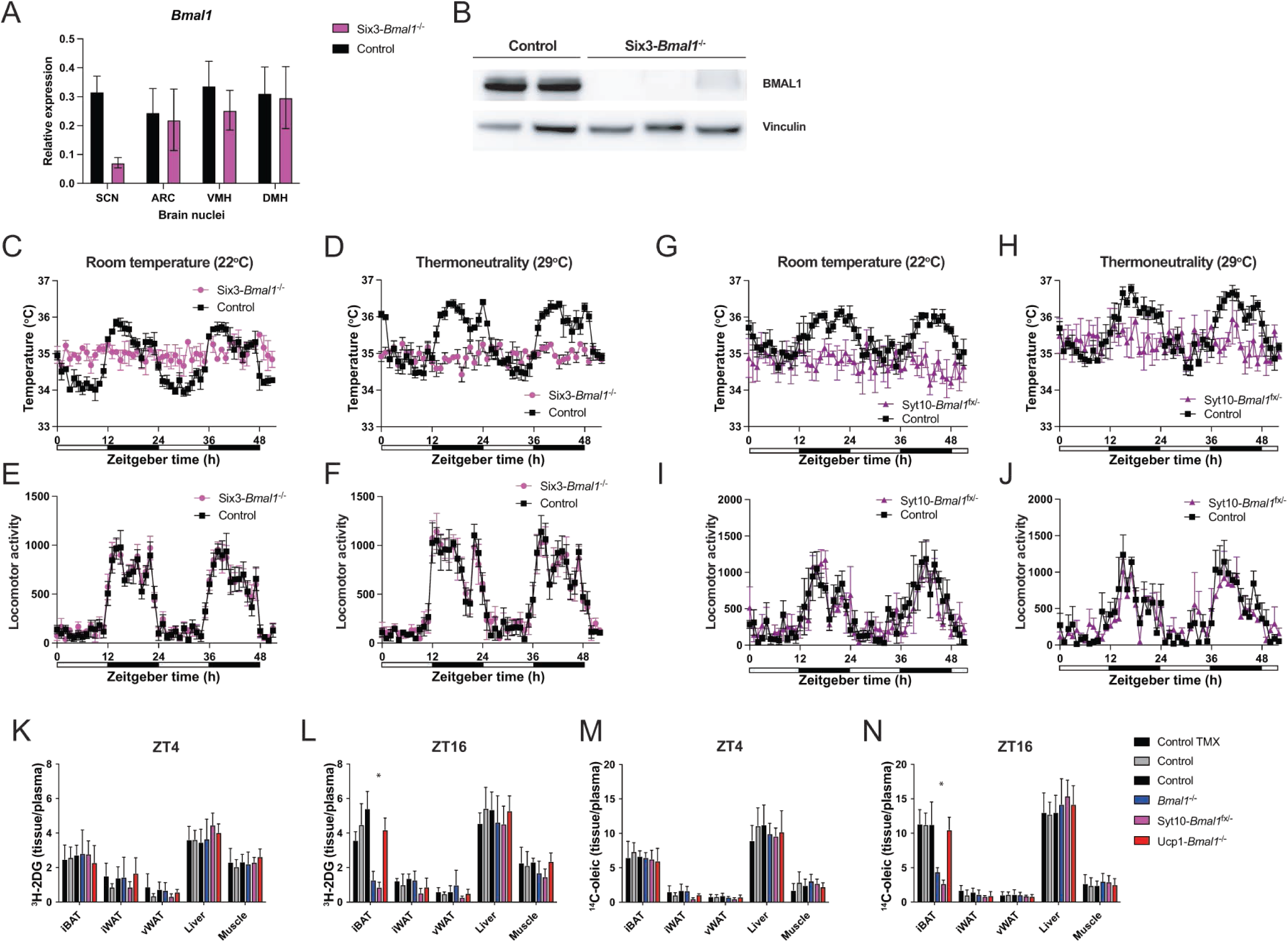

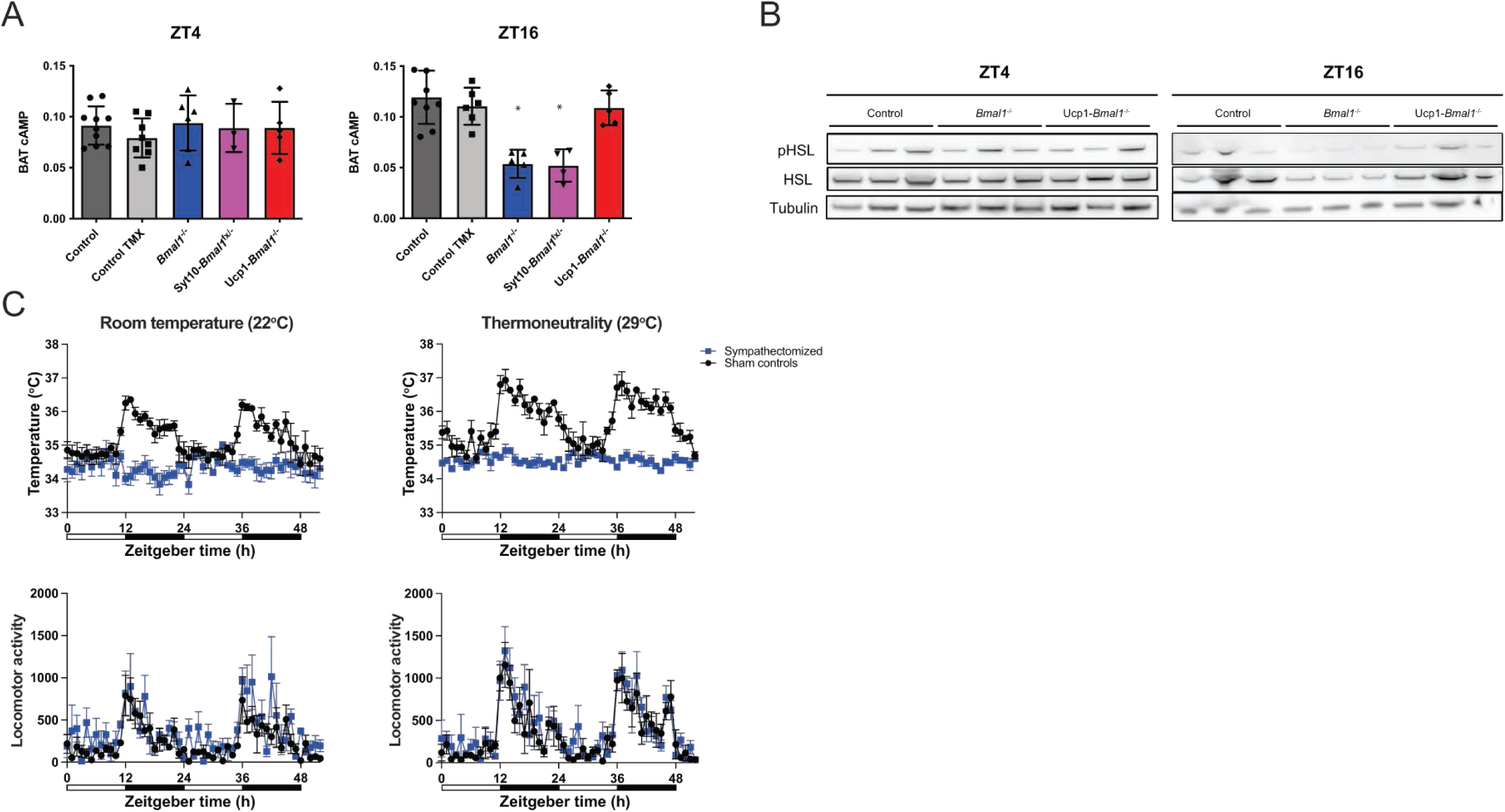

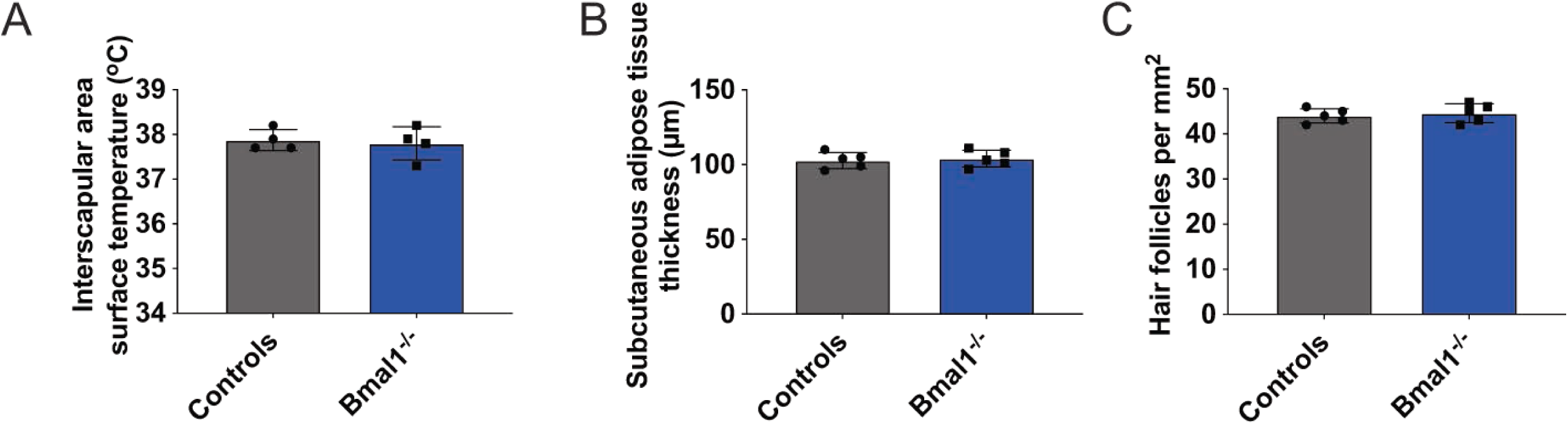

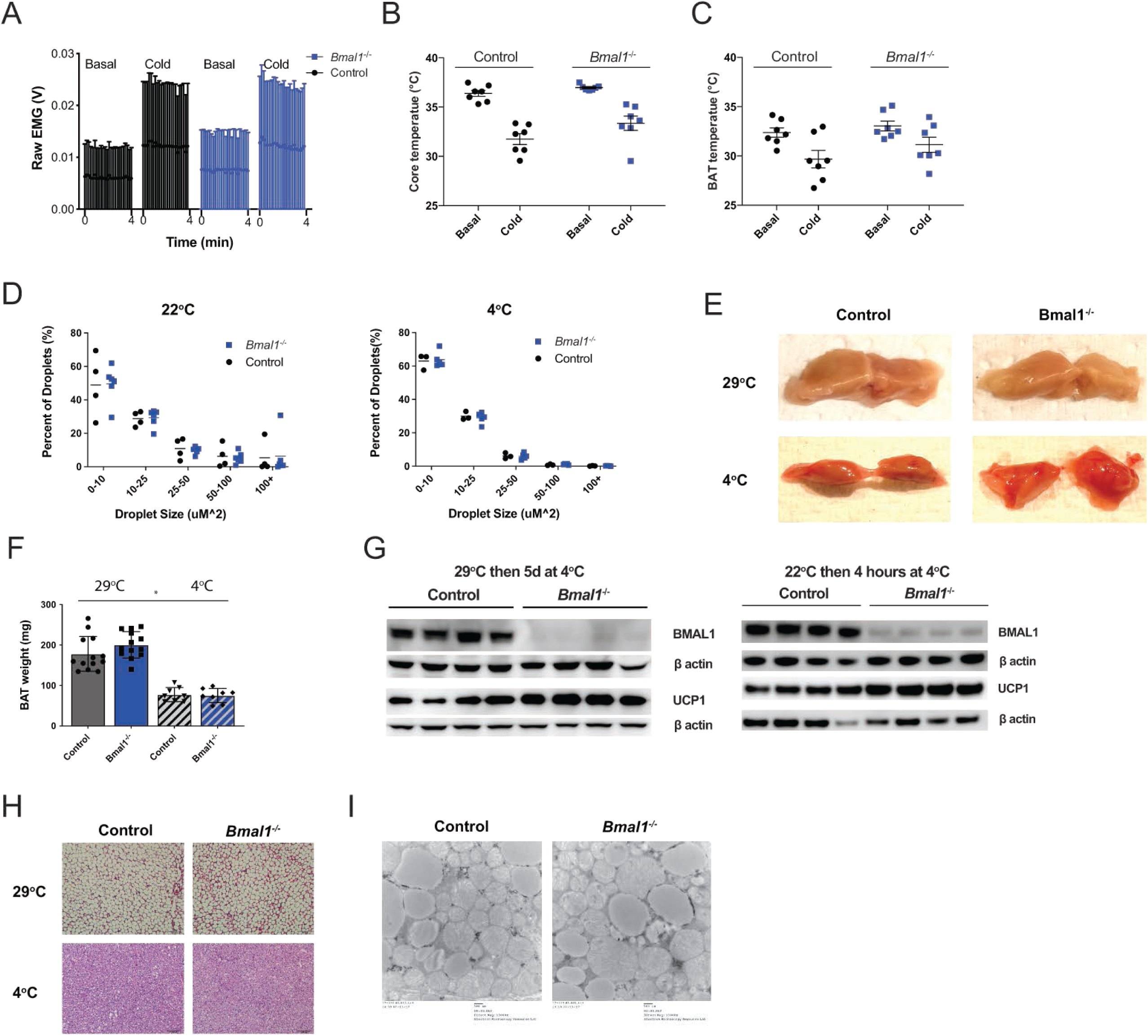

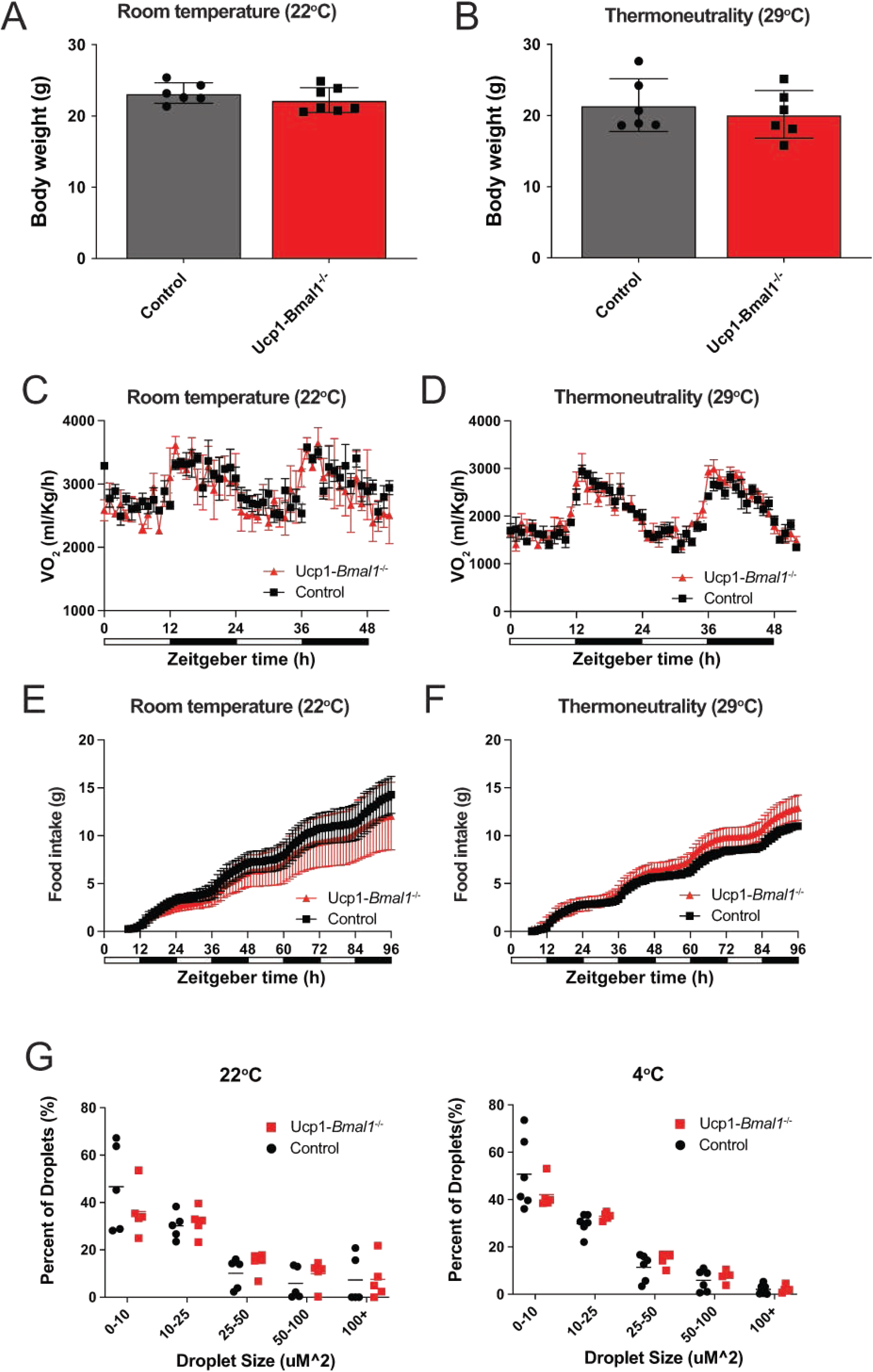

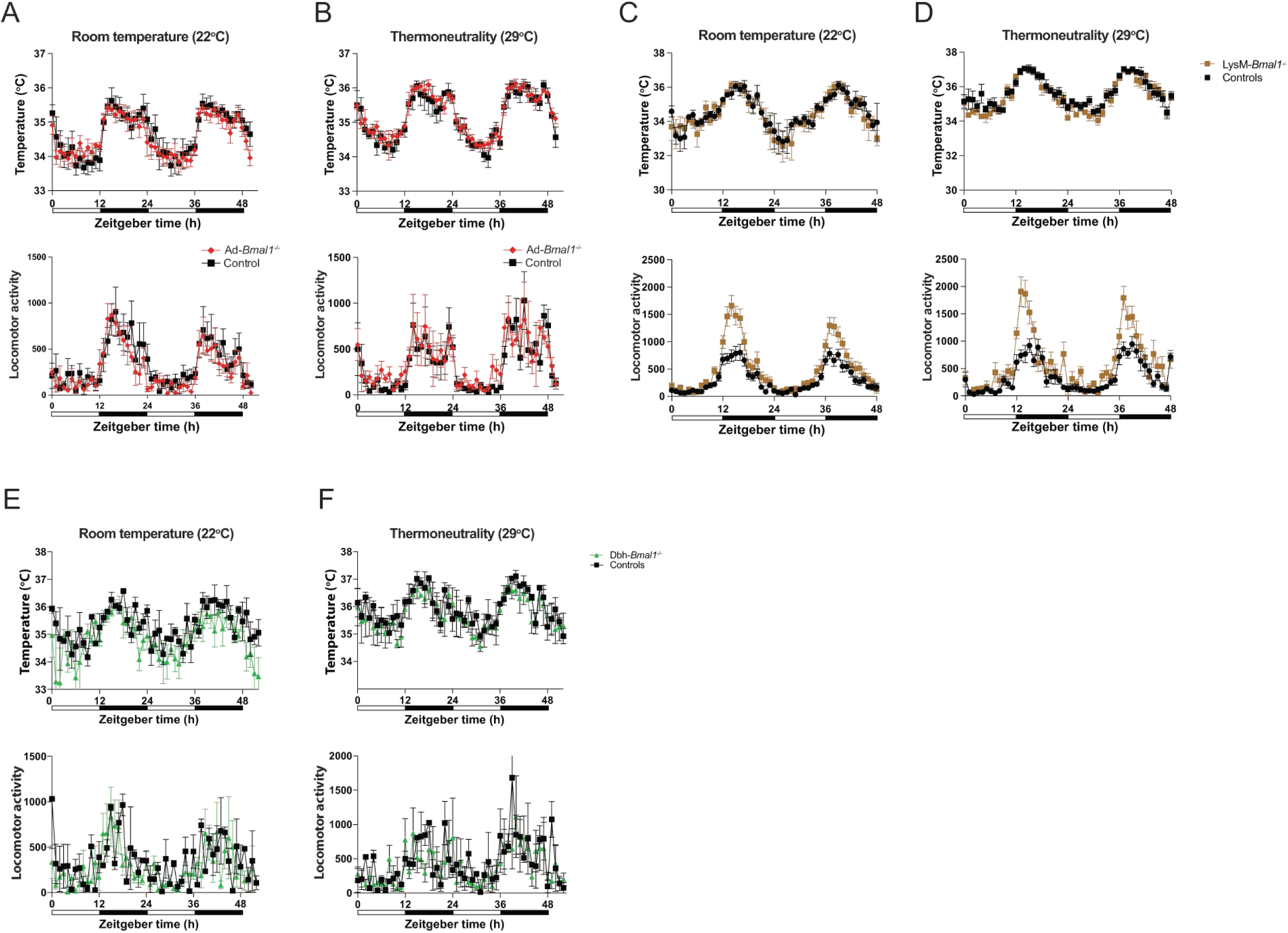

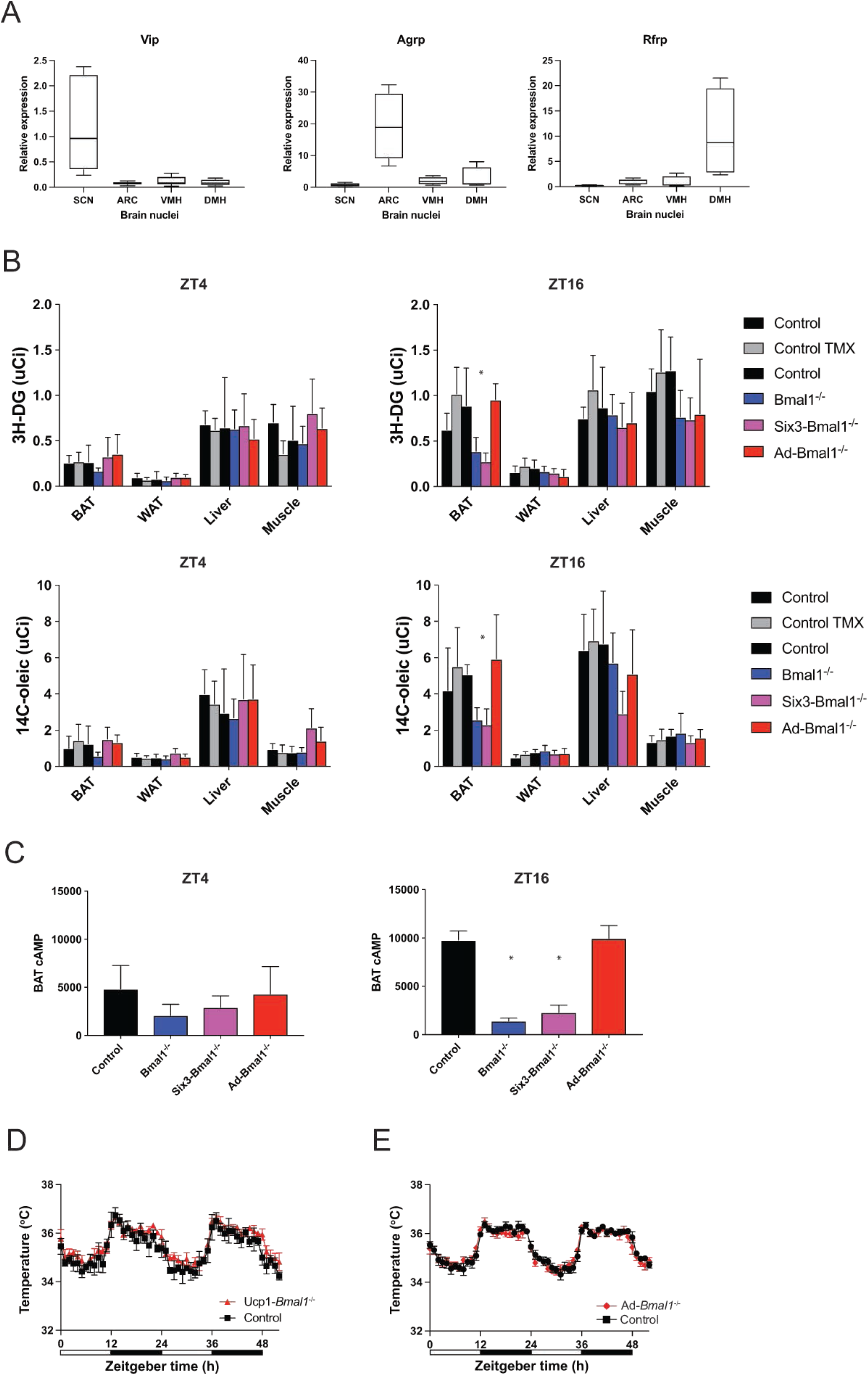

