## Supplemental Methods for "Brown adipose tissue thermogenesis rhythms are driven by the SCN independent of adipocyte clocks"

Supplementary Methods

Primers

| **Tbp** | 5'-ACCTTATGCTCAGGGCTTGG-3' | 5'-TGCCGTAAGGCATCATTGGA-3' |
| --- | --- | --- |
| **Vip** | 5'-GAACTTCAGCACCCTAGACAG-3' | 5'-GAAGAGTATCAGGAATGCCAGG-3' |
| **AgRP** | 5'-CAATGCCTTTTGCTACTGCC-3' | 5'-CCCATCCTTTATTCTCATCCCC-3' |
| **Rfrp** | 5'-GCTGGCAGATCATGACGTAGAG-3' | 5'-CACAGAAACCCCTGCACTCA-3' |
| **Bmal1** | 5’-AGGCCCACAGTCAGATTGAAA-3’ | 3’-CCAAAGAAGCCAATTCATCAATG-5’ |
